# An *in silico* map of the SARS-CoV-2 RNA Structurome

**DOI:** 10.1101/2020.04.17.045161

**Authors:** Ryan J. Andrews, Jake M. Peterson, Hafeez S. Haniff, Jonathan Chen, Christopher Williams, Maison Grefe, Matthew D. Disney, Walter N. Moss

## Abstract

SARS-CoV-2 is a positive-sense single-stranded RNA virus that has exploded throughout the global human population. This pandemic coronavirus strain has taken scientists and public health researchers by surprise and knowledge of its basic biology (e.g. structure/function relationships in its genomic, messenger and template RNAs) and modes for therapeutic intervention lag behind that of other human pathogens. In this report we used a recently-developed bioinformatics approach, ScanFold, to deduce the RNA structural landscape of the SARS-CoV-2 transcriptome. We recapitulate known elements of RNA structure and provide a model for the folding of an essential frameshift signal. Our results find that the SARS-CoV-2 is greatly enriched in unusually stable and likely evolutionarily ordered RNA structure, which provides a huge reservoir of potential drug targets for RNA-binding small molecules. Our results also predict regions that are accessible for intermolecular interactions, which can aid in the design of antisense therapeutics. All results are made available via a public database (the RNAStructuromeDB) where they may hopefully drive drug discovery efforts to inhibit SARS-CoV-2 pathogenesis.

## INTRODUCTION

There are four subfamilies within Coronaviridae – Alphacoronavirus, Betacoronavirus, Gammacoronavirus, and Deltacoronavirus. SARS-CoV 2 (severe acute respiratory syndrome coronavirus 2) falls under Betacoronavirus, with the subgenus Sarbecovirus^1^. The genome of SARS-CoV-2 is a roughly 30kb, positive sense (i.e. translation competent), 5′ capped single-stranded RNA molecule. The first two-thirds of Coronavirus genomes contain the ORF1ab gene, which consists of 16 <u>n</u>on-<u>s</u>tructural <u>p</u>roteins (NSP). ORF1ab consists of two large polyproteins (ORF1a and ORF1b) that are processed into smaller subunits. Translation begins in ORF1a, and following an RNA structural frame<u>s</u>hift element (FSE) translation resumes into the −1 frame of ORF1b; each are then processed into smaller units via proteases^2^. The latter third consists of the main viral structural elements (spike (S), envelope (E), membrane (M), and nucleocapsid (N)) as well as at least 13 known downstream ORFs. Upon infection of the host cell, viral RNA is translated into a series of NSP that replicate viral genomic (g)RNA and subgenomic (sg)RNA^2^. The replication-transcription complex (RTC) is capable of either continuous (genomic) or discontinuous (subgenomic) transcription into negative templates. Due to the method of template generation observed in SARS-like coronaviruses, negative sense templates share the same 5′ and 3′ end, differing in length based on their transcription regulatory sequences (TRS)^2^. TRS core sequences (AGCAAC or CUAAAC) have been previously observed to be highly conserved^3^ and facilitate addition of the nascent transcript to the 5′ leader TRS via complementary base pairing interactions between RNA structural elements. Both the 5′ leader and 3′ end sequences contain conserved RNA structural elements which are thought to serve function at both the genomic and subgenomic level. Viruses utilizing similar replication and expression strategies are known to contain a variety of RNA structural elements to direct these processes, however in the novel SARS-CoV-2 genome (and many other coronaviruses), the extent of functional RNA structural elements is still under investigation.

The recent global pandemic has put significant pressure on the academic community to provide structure and mechanism to the many unknowns associated with SARS-CoV-2. Particular focus is on the development of novel therapeutic agents to target and inhibit processes critical to SARS-CoV-2 infection and replication. To help address this important need, we have analyzed the SARS-CoV-2 genome using an RNA structural analysis and motif discovery pipeline, ScanFold, which has been previously applied to two other RNA viruses: Zika and HIV-1^4^; as well as human mRNAs for MYC^5^ and α-synuclein^6^. A significant innovation of ScanFold is that unique consensus secondary structural motifs are generated from bases pairs with the strongest evidence of being functional, while not necessarily focusing on evolutionary conservation (as in a recent analysis of SARS-CoV-2^7^). This provides a means of deducing novel structural elements that may only occur in this most recent pathogenic strain and, additionally, gives a global view of local structure that may not be forming specific structures – but still makes use of molecular stability. In addition to improving our basic understanding of RNA secondary structure, these results provided leads for the rational design of small-molecule drugs against SARS-CoV-2 RNAs^8^.

## RESULTS AND DISCUSSION

### SARS-CoV-2 ScanFold Analysis

Computational characterizations of RNA molecules of this size require special routines (typically a scanning analysis window) to perform the most accurate thermodynamic and statistical predictions of RNA folding. The program ScanFold was designed for this purpose and, in this study, has been applied to the reference sequence of SARS-CoV-2 (NC_045512.2); it has previously been used to characterize other positive-strand viral genomes including HIV-1 and Zika virus (ZIKV)^4^. The ScanFold method attempts to highlight regions most likely to have functional structure and generates unique 2D models for highly-structured and likely functional motifs^2^. This method is not only valuable for functional RNA structured motif discovery and mapping the general RNA folding landscape^9^, but for identifying structures likely to be available for targeting via small molecules as well^8,10^. The full output of these results can be seen in **Supplementary Dataset 1** and are available for browsing at https://www.structurome.bb.iastate.edu/sars-cov-2.

### ScanFold-Scan Analysis

Using ScanFold-Scan (performs initial scanning window analysis^1^), a variety of RNA folding metrics were calculated across the genome, which are described in detail in Methods section. Using a step size of 1 nt and a window size of 120 nt resulted in the generation of 29,783 analysis windows. The parameters used were previously found to perform optimally at detecting known structures in viral genomes that were consistent with experimental biochemical structure probing data^2^. The full SARS-CoV-2 scan results can be accessed at structurome.bb.iastate.edu/jbrowse where raw metrics are mapped directly alongside the genome sequence (and can be viewed in table format as well in **Table S1**).

Two of the most revealing metrics are shown in **Figure 1a**: the <u>m</u>inimum free <u>e</u>nergy (MFE) ΔG°, which reports the predicted change in the Gibb’s free energy for the most stable predicted RNA 2D structure and the ΔG° z-score, which measures the number of standard deviations more thermodynamically stable this MFE is vs. random sequences of the same nucleotide composition (Eq. 1; Methods). In this way, the ΔG° z-score attempts to characterize the propensity of a sequence to be *ordered* to adopt a particular fold, implying potentially functional roles for the structure (where negative z-scores are ordered to be more stable than expected for their composition). The MFE values across the genome ranged from −8.8 to −47.5 and averaged −26.1 kcal/mol (where MFE values mostly corresponded to changes in nucleotide content; **Table S1**). The ΔG° z-scores across the genome ranged from −6.4 to +2.74 and strikingly, yielded an average ΔG° z-score of −1.50. For comparison, this value is one standard deviation more stable on average than the previously scanned human pathogen positive strand RNA genomes of HIV-1 and ZIKV which had average ΔG° z-scores of −0.45 and −0.55 respectively. In each case, the z-scores are normally distributed around a negative median value (**Fig. 1b**) however, SARS-CoV-2 is sufficiently shifted into the negative (i.e. a median ΔG° z-score <−1) to be classified as having globally ordered RNA structure^11-13^. Local regions of highly negative z-scores and MFE values are found throughout the entirety of the genome and do not immediately appear concentrated in any given genomic region. However, two known RNA structural motifs annotated for human coronaviruses in Rfam^14,15^ overlapped regions predicted with negative z-scores: SL1-2 and the FSE (**Fig. 1c and 1d**).

**Figure 1.**
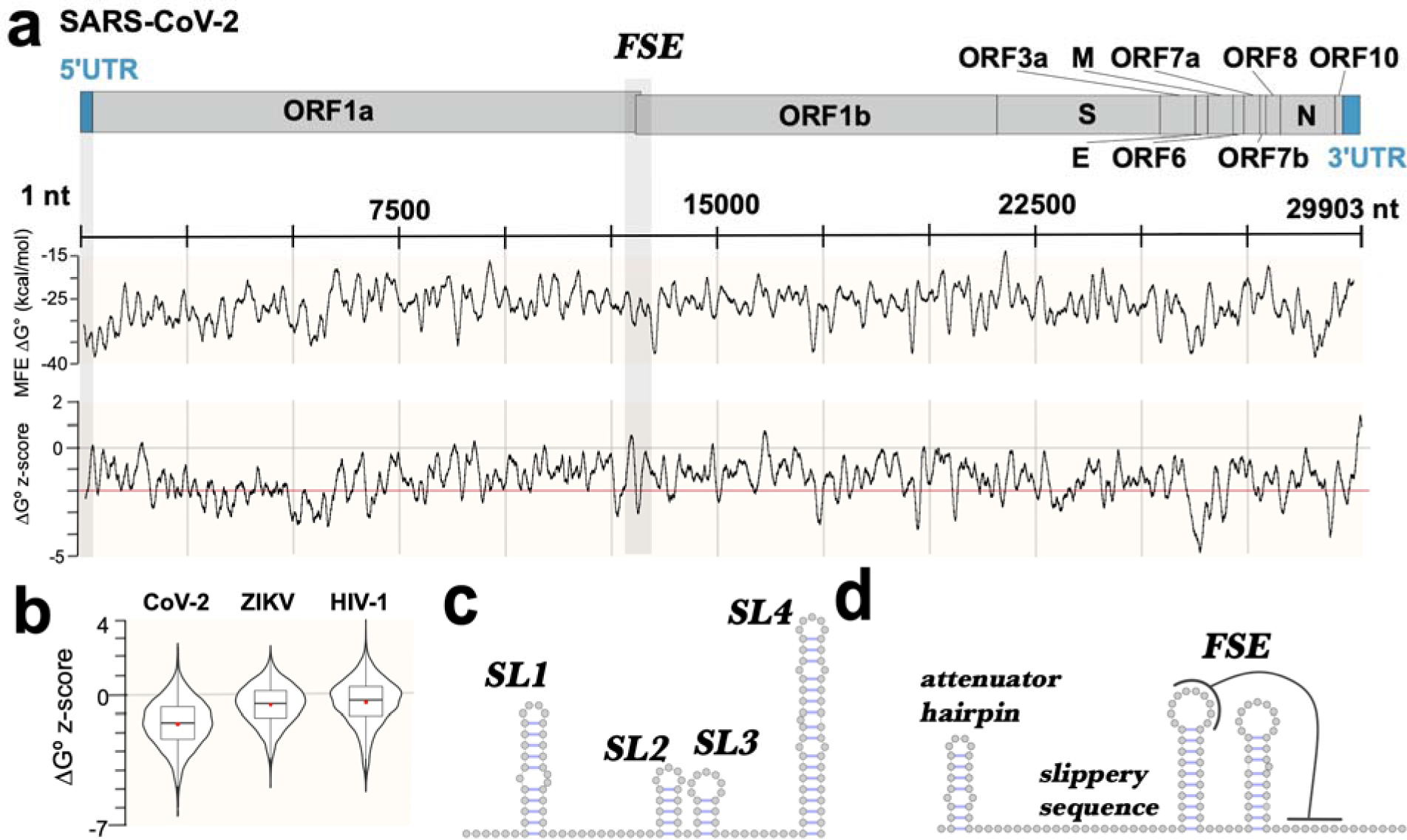
Overview of RNA folding analysis of the SARS-CoV-2 genome. **a)** Genomic features and coordinates are labeled at the top and annotated based on reference sequence NC_045512.2. The FSE element has been highlighted to show its location within two overlapping coding sequences in ORF1. The results of a ScanFold-Scan analysis (Methods) are shown below, mapped to the corresponding regions of the genome. The MFE and z-score are depicted using a 120 nt moving averages of values, raw values can be seen in Table S1. **b)** Overall distribution of raw z-score values calculated across the genome are shown alongside two other positive strand RNA genomes, ZIKV and HIV-1, which were analyzed using the same parameters as SARS-CoV-2. **c)** Generic model of the first four stem loops found in the 5′ UTR of *Betacoronavirus* genomes^7,18^(based on the Rfam entry for SL1-2; RF02910). **d)** Generic model showing the general architecture of the frameshift element (FSE) found in the similar coronavirus genomes^11^ (based on Rfam entry RF00507).

Positive ΔG° z-score regions can be seen throughout the genome as well, but are less frequent and smaller in size. Previous analyses found that such regions were more likely to be reactive to structure probing molecules^2^ (i.e. suggesting they are unstructured or highly dynamic)—potentially to facilitate intermolecular or long-range intra-genomic interactions. Of the 118 windows overlapping the start codon of *ORF1a*, 70 windows (i.e. 60%) had positive ΔG° z-scores (**Fig. 2a**) suggesting a preference for weak structures localizing around the start codon; consistent with previous analyses of RNA folding near start codons^5,16^. Another notable positive ΔG° z-score region is the 3′UTR, which was found to yield mostly positive z-scores; despite a higher than average GC content for this region (0.45 on average; **Table S1**), MFE values here were less stable than expected, averaging ΔG° z-scores of +0.98 (or roughly one standard deviation *less* stable than random). Scans were also performed in the negative sense of the genome, revealing slightly less propensity for structure. The MFE values ranged from −42.9 to −6.0 and averaged −23.25 kcal/mol; z-scores in the negative sense had a similar range of ΔG° z-scores as the positive sense (−5.76 to 2.66) but averaged lower at −1.12 (Table S1). Interestingly, ΔG° z-scores for the 3′ UTR in negative sense were *more skewed to the negative* (finding minimums as low as −2.32) resulting in an average z-score of 0.16 for the region.

**Figure 2.**
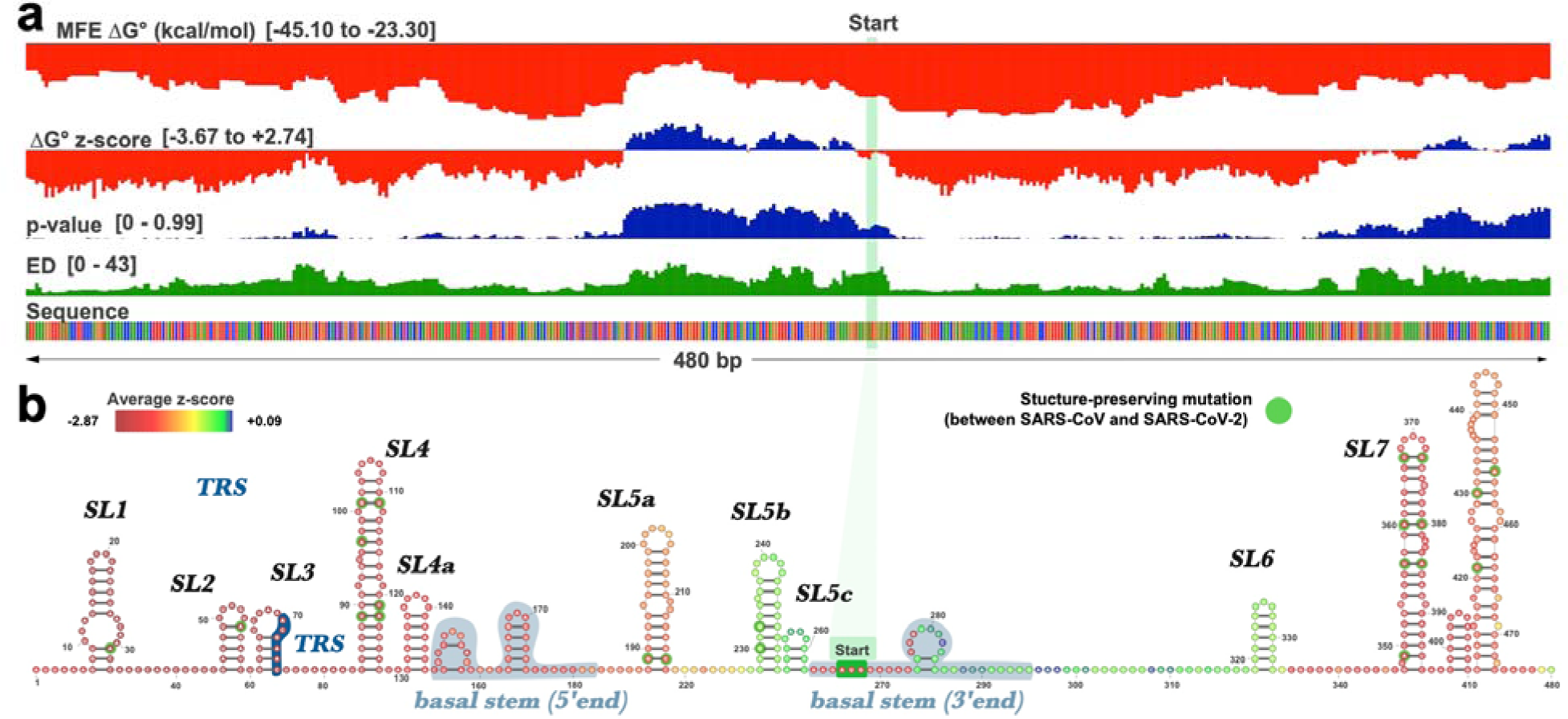
Full analysis of the 5′ UTR of SARS-CoV-2. **a)** The results of the full ScanFold pipeline are shown. ScanFold metrics and base pairs have been loaded into the IGV desktop browser^36^. Metric type and ranges are shown on the left side of the panel (metric descriptions can be found in methods). Here the start codon has been highlighted with a green bar and structures which correspond to previously named elements have been annotated. **b)** ScanFold RNA 2D structures are shown for the 5′UTR. All base pairs shown are consistent between SARS-CoV and SARS-CoV-2, and nucleotide variations which are present within structures have been highlighted with green circles. Structures have been visualized here using VARNA^37^.

### ScanFold-Fold Analysis

To predict the RNA secondary structures responsible for negative ΔG° z-score metrics, ScanFold-Fold was used to generate local RNA structural models based on the genome-wide ScanFold-Scan results. Here, each base pair observed throughout the scan is recorded along with corresponding metrics. Throughout the scan, each nucleotide was typically found to pair with multiple different partners as the scanning window stepped along the sequence; the ScanFold-Fold algorithm creates a comprehensive pair list (as well as all associated metrics) and highlights which base pairs *consistently* yielded low ΔG° z-scores^1^. This process is performed for each nucleotide, ultimately generating a single structural model. During the initial scan, 61,774 unique bp were observed (**Table S2**); utilizing the ScanFold-Fold algorithm to select only those base pairs with the most favorable ΔG° z-score metrics resulted in 7,235 bp (**Table S3**). Of these base pairs, 6,134 had average ΔG° z-scores less than or equal to −1 and, remarkably, 2,956 had average z-scores less than or equal to −2 (or around ∼41% of all base pairs, spanning 19.8% of the genome). This dramatic prevalence of ordered RNA folding is consistent with a recent study which found almost half the SARS-CoV-2 genome contained regions with conserved RNA structure^11^.

### Results in the 5′ UTR

The full ScanFold-Scan results for the 5′ UTR can be seen in **Figure 2a**. ScanFold-Fold modeled four of the known stem loops in the 5′ UTR leader region with z-scores < −2 (**Fig. 2b**). The start codon has been modeled as being unpaired, as opposed to previous global models which placed the start codon within a large multibranch structure (known as SL5; **Fig. S1**)^7,17,18^. As reported above, the scanning data around the start codon resulted in positive ΔG° z-scores, which in this case favor the formation of a small hairpin where the 5′ end of the SL5 basal stem would form and keeps the start codon nucleotides unpaired (**Fig. 2b**). However, since the basal stem base pairs span >120 nt (the window size used), we would not expect ScanFold to identify it. The ScanFold model leaves 75% of the basal stem nucleotides unpaired, indicating that local folds may not strongly compete against formation of the larger stem. Further, though the basal stem of SL5 is not present in the ScanFold model, the terminal stem loops (SL5a-c) *are* modeled consistent with recent global models of SL5^7,18^ (**Fig. 2b)**.

### Frameshift Element

The FSE is an RNA structural motif which incorporates nucleotides of the overlapping frames of ORF1a and ORF1b (nt 13476 to 13542; **Fig. 3a**). The full ScanFold-Scan results for the FSE region can be seen in **Figure 3a**, where the 120 nt windows fully incorporating those nucleotides (i.e. windows around nt 13,420, or 120 nt upstream of the elements end) were found to yield the most negative z-scores. The base pairs which correspond to these negative values are shown in the ScanFold-Fold model (**Fig. 3b**). The ScanFold-Fold model of the FSE is largely consistent with recent models^7,19^. This model consists of two stable hairpins: the first of which contains a loop sequence which forms a pseudoknot by pairing with nucleotides upstream of the second hairpin (**Fig. 3b**). ScanFold cannot predict the pseudoknot directly, however the generated model does leave the pseudoknot forming nucleotides sufficiently unpaired to allow for the interaction to occur. Functional elements upstream of these hairpins are placed into an alternative model by ScanFold. Here, the attenuator hairpin is embedded in a multibranched structure along with the slippery sequence, which is predicted to form a small three base pair stem. Notably, the only bps which had average z-scores < −2 for the FSE region are found in the basal stem of this previously unreported multibranched structure. These findings suggest the full frameshift element may incorporate more nucleotides than previously described.

**Figure 3.**
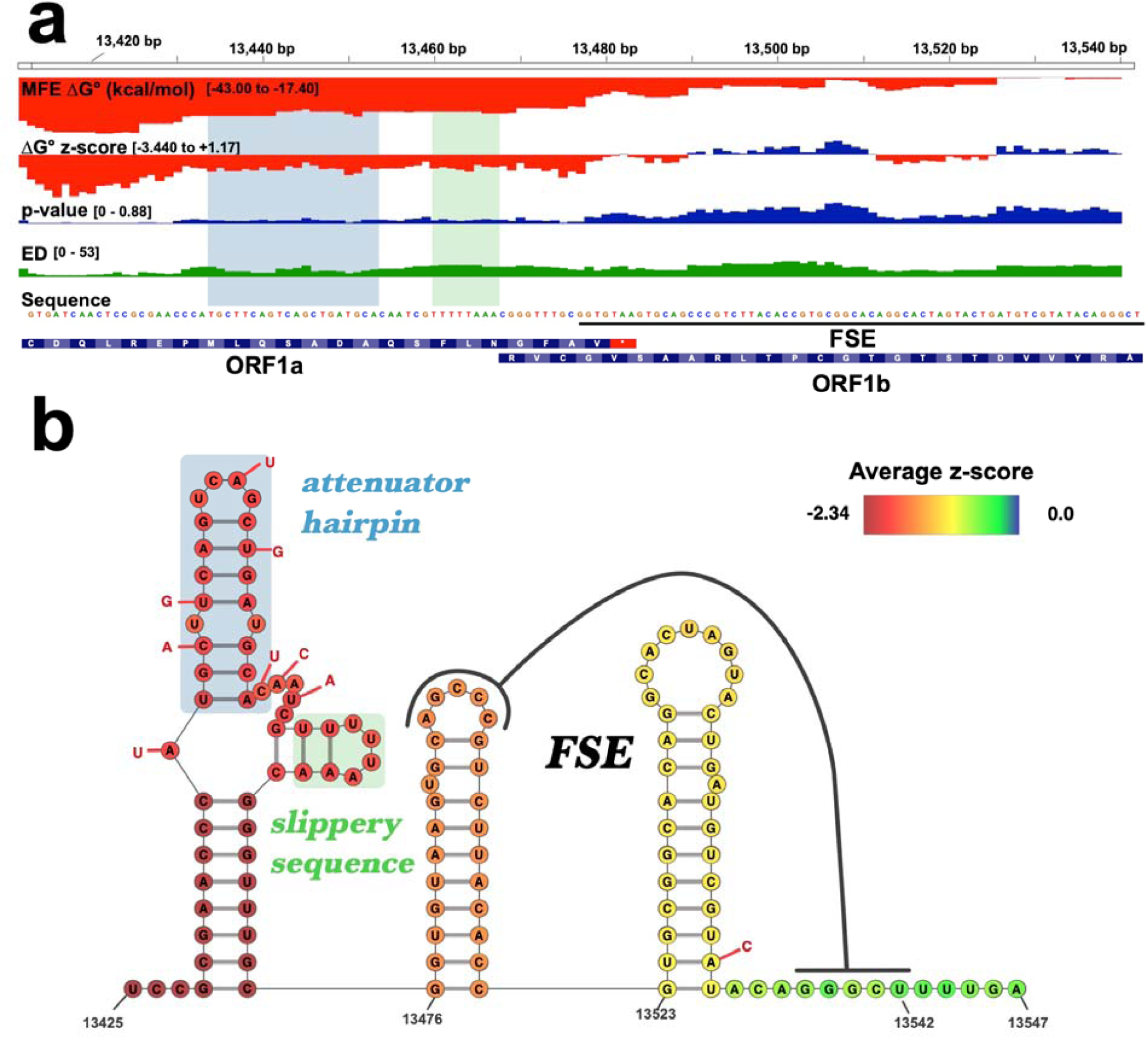
Full analysis of the FSE region from SARS-CoV-2. **a)** The results of the full ScanFold pipeline are shown. ScanFold metrics and base pairs have been loaded into the IGV desktop browser^36^. Metric type and ranges are listed on the left side of the panel (metric descriptions can be found in the Methods section). Arc diagrams for ScanFold model base pairs are colored based on their average z-score (blue for <= −2, green for <= −1, and yellow for <= 0). **b)** The ScanFold RNA 2D structures for the full FSE region are shown. All differences between the SARS-CoV and SARS-CoV-2 primary structure are reported directly next to the corresponding nucleotide of SARS-CoV-2 (with the corresponding SARS-CoV nucleotide in a red circle). Structures have been visualized here using VARNA^37^.

### Results in the 3′ UTR

The 3′UTRs of the *Sarbecovirus* genomes contain two RNA structural elements; a 3′ UTR pseudoknot structure presumably required for replication^20^ and a mobile genetic element with an undetermined function known as the 3′ stem-loop II-like motif (s2m)^21^. Under the current genome annotation (NC_045512.2) much of the previous 3′ UTR sequence is now found within an upstream open reading frame named ORF10 (however this has been recently reported as being an untranslated ORF as well^22^). ScanFold’s model partially recapitulates a recent model of the region^7^ (blue pairs; **Fig. S2a**), however, overall metrics for the region (including high ensemble diversity values and positive ΔG° z-scores; **Table S1**) suggest the region is unstructured and/or highly dynamic. As such, the ScanFold model predicts the downstream region to be mostly absent of structured elements. However, as mentioned earlier, the 3′ UTR in the negative sense actually yielded several significantly low z-score windows. Here, ScanFold-Fold predicts this signal to arise from a large internally looped hairpin whose terminal stem consists of the reverse complemented nucleotides of the s2m element (**Fig. S2b**). Notably, the internal loop of the negative strand 3′ UTR structure, as in the forward strand, consists of a deeply conserved eight nucleotide primary sequence motif ^17^. The unpaired nucleotides opposite this sequence are predicted to have highly negative z-scores, suggesting flanking nucleotides may be ordered to encourage internal loop formation (**Fig. S2b**). This structure in many ways mirrors the structure reported for the positive sense (**Fig. S2**) which is not uncommon for RNA structured elements^23^. In the negative strand however, the structure is more thermodynamically stable yielding a more negative z-score – properties which provide evidence that RNA structure may play a functional role in the negative strand as well^23^.

### Novel Motifs with Strong Predicted Metrics for Structure/Function

Beyond recapitulating elements of described structure within the 5′ UTR, FSE, and 3′ UTR, ScanFold-Fold predicted over 500 individual motifs with strong predicted metrics for structure/function. All motif models can be accessed on the RNAStructuromeDB (https://structurome.bb.iastate.edu/download/sars-cov-2-extractedstructures) and in Table S4. Many motifs have better prediction metrics than either of the previously described motifs. For example, the most negative ΔG° z-scores are found downstream of ORF1b (**Fig. 1a**). Notably, this region of the genome is discontinuously transcribed into negative-sense RNA fragments which serve as intermediates during the generation of positive sense sgRNAs. Although the mechanism of discontinuous transcription is currently unknown, structural elements are known to play a functional role in terminating negative strand transcription and/or generating sgRNA in other viral genomes.

There are seven core TRS sequences (5′-ACGAAC-3′) present in the genome. These are found between sgRNA coding regions and are involved in the addition of the 5′ leader region of the genome onto each sgRNA. The mechanism is still unclear, however, current models suggest leader addition is directed by base pairing between nascent negative strand TRS sequences to the positive strand leader TRS^22^. All of the TRS core sequences are present in ScanFold predicted motifs (Table S4). The region encompassing the E coding sequence (E_CDS_) contains two of these TRS sequences in relatively close proximity (the E_CDS_ is a small ORF spanning only 228 nt). ScanFold results for this region are shown in **Figure 4**. Notably, the start codon here is overlapped by negative z-score windows (**Fig. 4a**), with individual windows reaching as low as −5. The ScanFold-Fold predicted model for the region finds these low z-scores generated via two hairpins near the start codon. The first hairpin incorporates both the TRS sequence and the start codon into its loop and the second hairpin is a stable structure, with average z-scores as reaching as low as −3.88 (**Fig. 4b**). Interestingly, each of these structures are predicted to resemble transcription terminator hairpins^24-26^, which could suggest a potential functional role.

**Figure 4.**
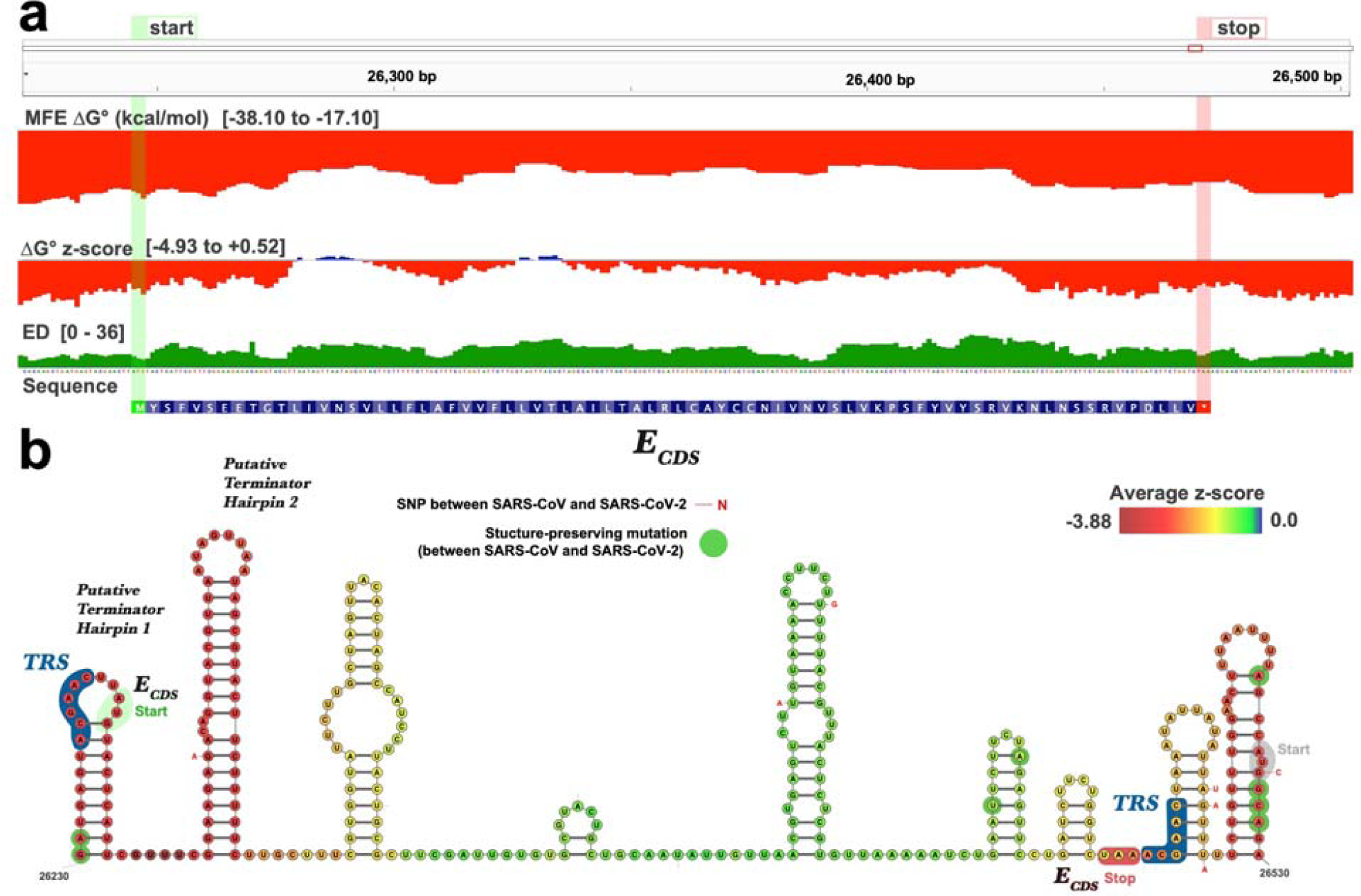
ScanFold results of the region encompassing the E_cds_. **a)** The full ScanFold-Scan metrics are shown. **b)** ScanFold-Fold predicted model for the region is shown here. Nucleotides here are colored according to their average z-score from Table S2. The TRS nucleotides have been highlighted and labeled with a dark blue outline, and the start and stop codons have been highlighted and labeled green and red respectively. Differences between SARS-CoV and SARS-CoV-2 have been annotated as described in Figures 2 and 3.

Immediately upstream of E_cds_ is a region in the ORF3a_cds_ which generated the lowest z-score windows of the genome (as low as −6.4). This is presumably due to a stretch of 10 predicted hairpins which each generate z-scores < −2 (when folded individually; Table S4). ScanFold finds this amount of ordering highly significant and models the 3′ end of the coding sequence to be mostly bound up in stable hairpins (**Fig. S3**).

### Conservation of ScanFold Predicted Structural Elements

To determine the prevalence of ScanFold predicted RNA structures between pathogenic coronavirus strains, a structural conservation analysis was performed between SARS-Cov, MERS and SARS-CoV-2. Recent analyses indicated the MERS genome was decidedly divergent from SARS-CoV-2, with only ∼50% pairwise identity between the genomes^27^ and too distant to yield powerful structural covariation support for conserved structure^7^. However, the pathogenic SARS genomes are much less distant, residing in the same *Sarbecovirus* genus of the *Coronaviridae* family^27^; suggesting a greater likelihood of harboring identical RNA secondary structural motifs. In order to determine the structural conservation of ScanFold models between SARS-CoV and SARS-CoV-2, an alignment was generated using MAFFT^28,29^. The alignment was consistent with earlier analyses resulting in a 79.0% pairwise identity for the primary structure with similar results for the secondary structure, finding 4,860 of the 6,134 bp with average z-scores <= −1 being completely conserved (79%). Mutations between the genomes were present in 2796 of ScanFold predicted base pairs with 1522 of those consistent with the predicted base pair and 1274 mutations inconsistent. While the overall secondary structures for each genome correlate mostly with primary structure, local regions were found to have higher degrees of conservation.

Consistent with their functional conservation across several *Coronaviridae* genomes^30,31^, the FSE and 5′ UTR regions had higher base pair conservation on average. The 5′ UTR stem loops (SL1-4 and SL5a-c) base pairs are 100% conserved between SARS-CoV and SARS-CoV-2, and when mutations were present, they were consistent with the ScanFold model (**Fig. 2b**). The two hairpins of the FSE had 20 of 21 base pairs conserved between SARS-CoV and SARS-CoV-2 with a single mutation present in the base of the second stem (**Fig. 3b**). The multibranch structure upstream, consisting of the attenuator and slippery sequence however, contained 8 mutations, 4 of which are directly within, and inconsistent with the SARS-CoV attenuator hairpin secondary structure (**Fig. 3b)**. This provides little evidence for conservation of specific base pairs in the attenuator, but rather, preservation of hairpin structures at these sites in both viruses. The novel base pairs which extend the FSE model were conserved between both genomes suggesting the basic architecture of this proposed extended structure could be present and functional in both genomes.

The ScanFold predicted structures corresponding to previously observed structural elements in the 3′ UTR (blue pairs; **Fig. S2a**) are completely conserved between SARS-CoV and SARS-CoV-2. The corresponding structured motifs detected in the negative strand are similarly conserved between SARS-CoV and SARS-CoV-2 (**Fig. S2b**). The negative strand motif consisting of the large internal loop, is conserved between genomes as well, but as in the positive strand, mutations do occur within the s2m region.

The novel motifs predicted near the E_cds_ TRS sequences have some evidence of conservation (with two consistent mutations found in the stem of the first putative terminator hairpin, and four consistent mutations in the final hairpin of the model containing the start codon of the M coding region; **Fig. 4b**). The structures found in the ORF3a region (**Fig. S3**) do not show evidence of specific base pair conservation, however, the same region in SARS-CoV does appear similarly structured, yielding z-scores as low as −5.49 (data not shown). This homologous region in SARS-CoV however is only ∼68% similar at the primary structure level and utilizes a different gene architecture (with two overlapping reading frames).

## METHODS

### In silico analyses

The SARS-CoV-2 (NC_045512.2) genome sequence was downloaded from the NCBI nucleotide database. ScanFold-Scan was run using a 120 nt window moving with a single nucleotide step size. Each window was analyzed using the RNAfold algorithm implemented in the ViennaRNA package (2.4.14)^32^. For each window the MFE ΔG° structure and value was predicted using the Turner energy model^33,34^ at 37°C. To characterize the MFE, a ΔG° z-score is calculated for each. Each MFE predicted for the native sequence (MFE_native_) is compared to MFE values calculated for 100 shuffled version of the sequence with the same nucleotide composition (MFE_random_) as shown in Eq. 1; using an approach adapted from Clote et. al.^35^ Here, the standard deviation (σ) is calculated across all MFE values. 

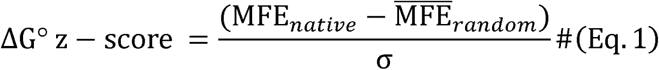

The p-value corresponds to the number of MFE_random_ values which were *more* stable (more negative) than the MFE_native_. In addition to these metrics, RNAfold partition function calculations are utilized to characterize the potential structural diversity of the native sequence. These include the <u>e</u>nsemble <u>d</u>iversity (ED) and the centroid structure. The centroid structure depicts the base pairs which were “most common” (i.e. had the minimal base pair distance) between all the Boltzmann-ensemble conformations predicted for the native sequence. The ED then attempts to quantify the variety of different structures which were present in the ensemble (where higher numbers indicate multiple structures unique from the predicted MFE and low numbers indicate the presence of a dominant MFE structure highly represented in the ensemble).

For comparisons made between the *Sarbecovirus* genomes, the SARS-CoV genome (NC_004718.3) was aligned to SARS-CoV-2 (NC_045512.2) using the MAFFT webserver^28^ with default settings (in this case the FFT-ns-i method^13^ was implemented).

## CONCLUSION

This work lays out the predicted local RNA folding landscape of the SARS-CoV-2 transcriptome. In addition to general trends in RNA structure, we present unique motifs from base pairs that contribute most to the exceptional thermodynamic stability of SARS-CoV-2 RNAs. This work lays out the potential druggable RNA structurome of SARS-CoV-2 to drive forward both basic research into structure/function as well as efforts to target this virus using small molecule therapeutics.

## ACKNOWLEDGEMENTS

This research was supported by NIH/NIGMS grants R00GM112877, R01GM133810, and by startup funds from the Roy J. Carver Charitable Trust to WNM; as well as grant R01-GM097455 to MDD.

---

**Figure S1.**
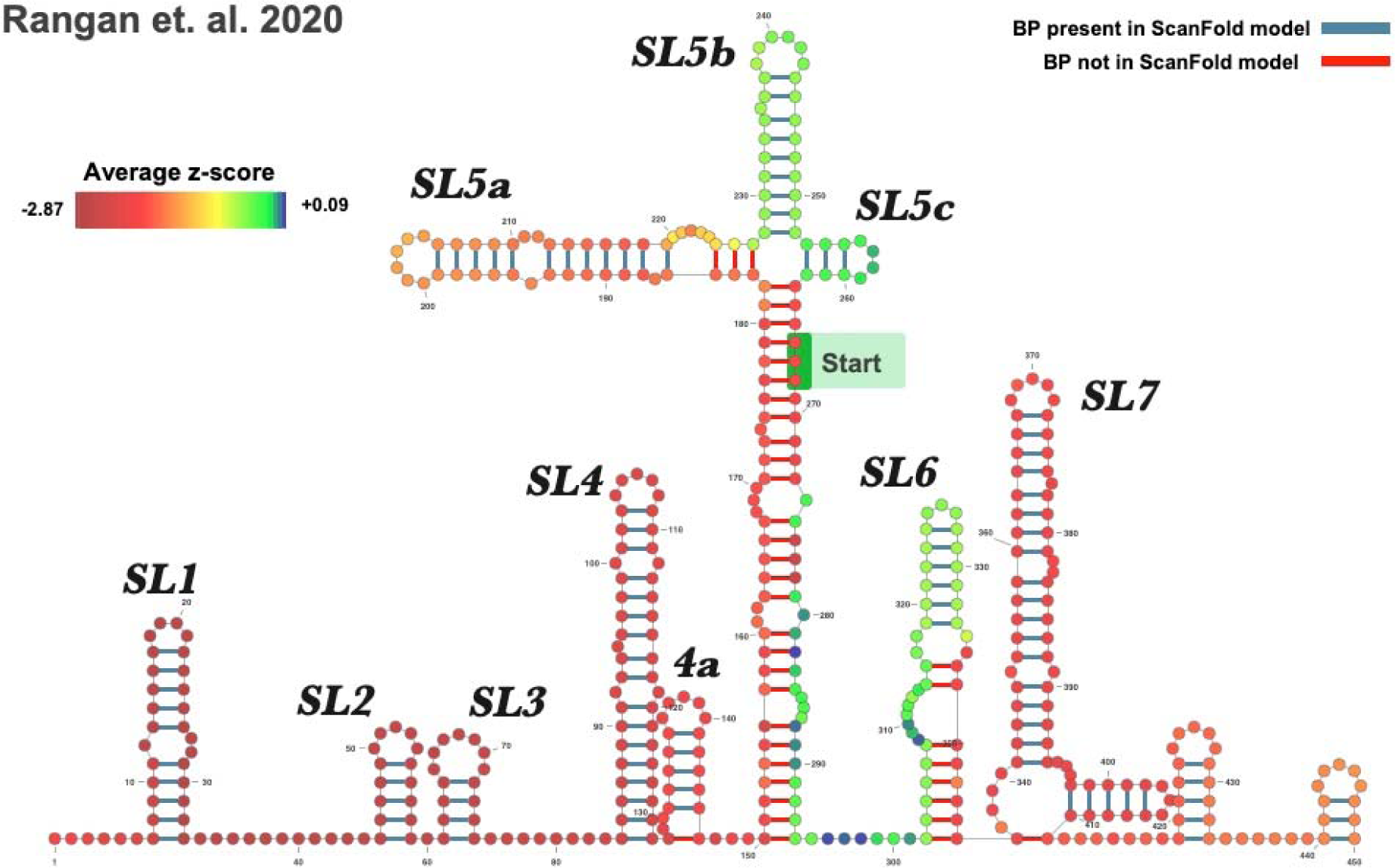
Recent global model of the 5′ UTR of SARS-CoV-2. Structure is shown as reported in Rangan et. al.^7^ Base pairs here are colored based on their presence in the ScanFold model from Figure 2, and nucleotides are colored based on the average z-score from Table S2.

**Figure S2.**
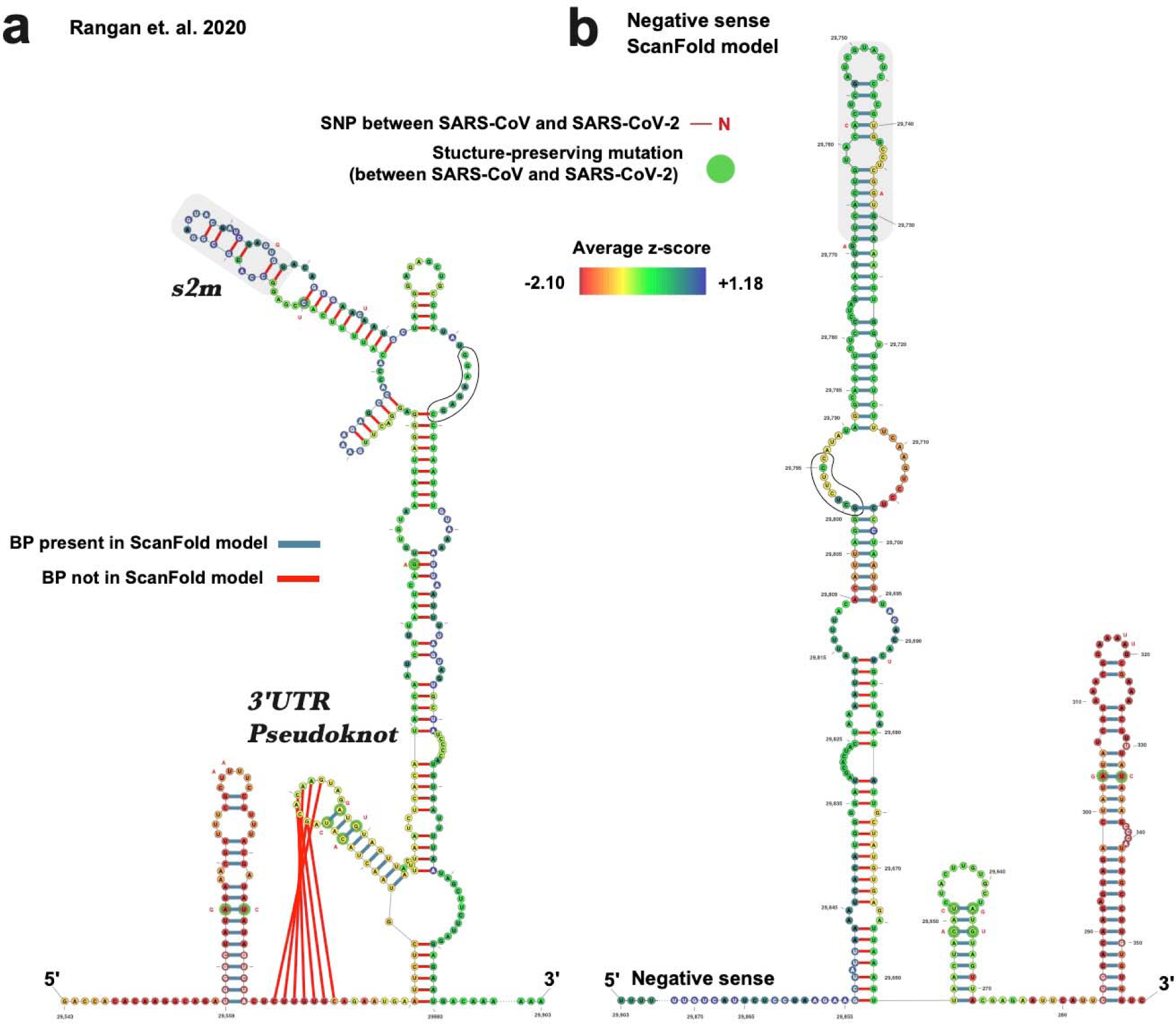
Model of the forward and reverse strand of the 3′ UTR of SARS-CoV-2. a) Model of the SARS-CoV-2 3′ UTR from Rangan et. al. annotated with ScanFold metrics. b) ScanFold model of the negative strand of the 3′UTR. Here the region was refolded in RNAfold using base pairs with z-score averages <-2 as constraints. Base pairs here are colored based on their presence in the ScanFold model of the region, and nucleotides are colored based on the average z-score from Table S2.

**Figure S3.**
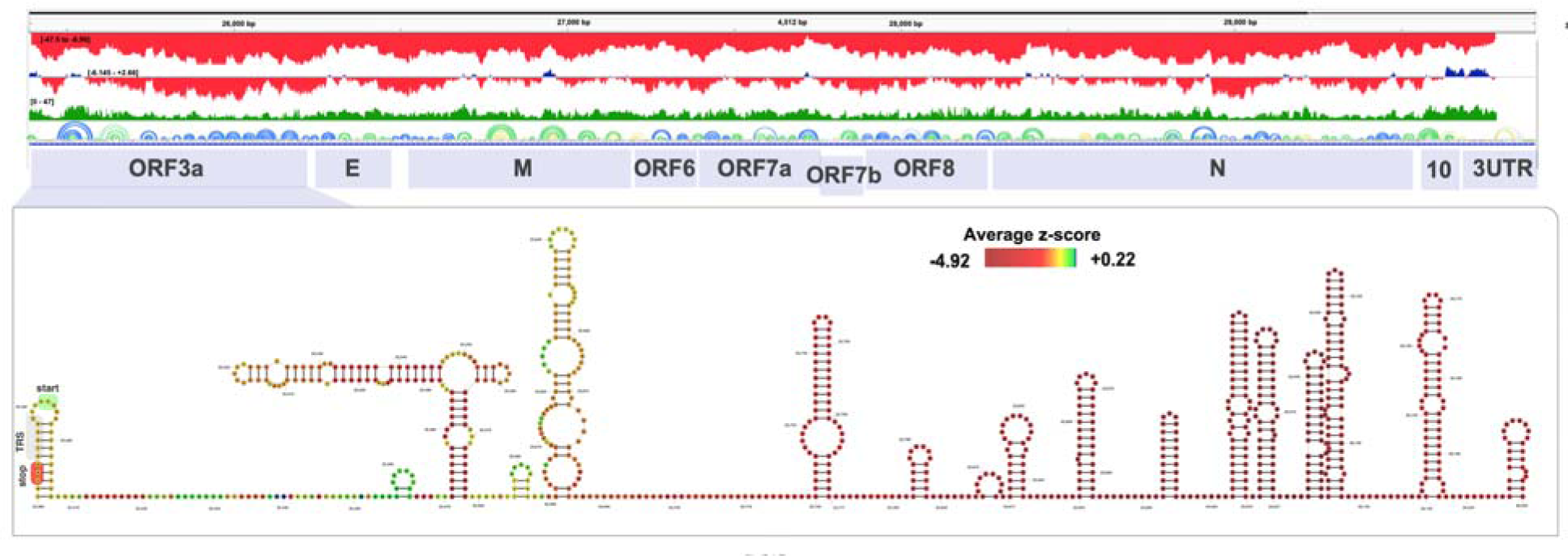
Model for the highly structured ORF3a coding sequence.

## Notes

https://www.structurome.bb.iastate.edu/sars-cov-2

### Competing Interest Statement

The authors have declared no competing interest.

